# Visualization and regulation of translocons in *Yersinia* type III protein secretion machines during host cell infection

**DOI:** 10.1101/431908

**Authors:** Theresa Nauth, Franziska Huschka, Michaela Schweizer, Jens B. Bosse, Andreas Diepold, Antonio V. Failla, Anika Steffen, Theresia Stradal, Manuel Wolters, Martin Aepfelbacher

**Affiliations:** Institute of Medical Microbiology, Virology and Hygiene, University Medical Center Hamburg-Eppendorf (UKE), Hamburg, Germany; Center for Molecular Neurobiology (ZMNH), University Medical Center Hamburg-Eppendorf (UKE), Hamburg, Germany; Heinrich Pette Institute, Leibniz Institute for Experimental Virology (HPI), Hamburg, Germany; Department of Ecophysiology, Max Planck Institute for Terrestial Microbiology, Marburg, Germany; UKE Microscopy Imaging Facility, University Medical Center Hamburg-Eppendorf (UKE), Hamburg, Germany; Department of Cell Biology, Helmholtz Centre for Infection Research (HZI), Braunschweig, Germany

**Author notes:** These two groups of authors contributed equally to this work. The authors have declared that no competing interests exist. Conceived and designed experiments: TN FH MW MA; Performed experiments: TN FH MW MS; Analyzed the data: TN FH MW JB; Contributed reagents/materials/analysis tools/microscopes: MS AS VF TS JB AD; Wrote the paper: MA TN MW.

## Abstract

Type III secretion systems (T3SSs) are essential virulence factors of numerous bacterial pathogens. Upon host cell contact the T3SS machinery - also named injectisome - assembles a pore complex/translocon within host cell membranes that serves as an entry gate for the bacterial effectors. Whether and how translocons are physically connected to injectisome needles, whether their phenotype is related to the level of effector translocation and which target cell factors trigger their assembly have remained unclear. We employed the superresolution fluorescence microscopy techniques Stimulated Emission Depletion (STED) and Structured Illumination Microscopy (SIM) as well as immunogold electron microscopy to visualize *Y. enterocolitica* translocons during infection of different target cell types. Thereby we were able to resolve translocon and needle complex proteins within the same injectisomes and demonstrate that these fully assembled injectisomes are generated in a prevacuole, a PI(4,5)P2 enriched host cell compartment inaccessible to large extracellular proteins like antibodies. Furthermore, the putatively operable translocons were produced by the yersiniae to a much larger degree in macrophages (up to 25% of bacteria) than in HeLa cells (2% of bacteria). However, when the Rho GTPase Rac1 was activated in the HeLa cells, uptake of the yersiniae into the prevacuole, translocon formation and effector translocation were strongly enhanced reaching the same levels as in macrophages. Our findings indicate that operable T3SS translocons can be visualized as part of fully assembled injectisomes with superresolution fluorescence microscopy techniques. By using this technology we provide novel information about the spatiotemporal organisation of T3SS translocons and their regulation by host cell factors.

**Author Summary:** Many human, animal and plant pathogenic bacteria employ a molecular machine termed injectisome to inject their toxins into host cells. Because injectisomes are crucial for these bacteria’s infectious potential they have been considered as targets for antiinfective drugs. Injectisomes are highly similar between the different bacterial pathogens and most of their overall structure is well established at the molecular level. However, only little information is available for a central part of the injectisome named the translocon. This pore-like assembly integrates into host cell membranes and thereby serves as an entry gate for the bacterial toxins. We used state of the art fluorescence microscopy to watch translocons of the diarrheagenic pathogen *Yersinia enterocolitica* during infection of human host cells. Thereby we could for the first time - with fluorescence microscopy - visualize translocons connected to other parts of the injectisome. Furthermore, because translocons mark functional injectisomes we could obtain evidence that injectisomes only become active when the bacteria are almost completely enclosed by host cells. These findings provide a novel view on the organisation and regulation of bacterial translocons and may thus open up new strategies to block the function of infectious bacteria.

## Introduction

Bacterial type III secretion systems (T3SSs) are molecular machines also termed injectisomes that translocate proteins of bacterial origin (i.e. effectors) into host cells. T3SSs are essential virulence factors of numerous human, animal and plant pathogens including *Chlamydia, Pseudomonas*, EPEC and EHEC, *Salmonella, Shigella* and *Yersinia* [2, 3]. Based on sequence identity among structural components nine T3SS families were classified [4], Whereas assembly, structure and function of the T3SSs are highly conserved, the biochemical activities of the translocated effectors often are multifaceted and reflect the infection strategies of the individual pathogens [5]. Because of their uniqueness in bacteria on one hand and central role for bacterial pathogenicity on the other hand T3SSs have been considered as targets for novel antiinfective strategies [6–9]. In addition, the ability of T3SSs to inject arbitrary proteins into immune cells has been exploited for experimental vaccination strategies [10].

Based on topology and function injectisomes can be separated into different parts: i) the sorting platform on the cytoplasmic side of the injectisome is a protein assembly thought to control targeting and secretion of the T3SS substrates; ii) the basal body including the export apparatus spans the inner and outer bacterial membranes; iii) the 30-70 nm long needle filament is built by a single multimerized protein and together with the basal body forms the needle complex; iv) the tip complex consists of a hydrophilic protein that caps the needle filament and regulates assembly of the translocon; v) the translocon consists of two hydrophobic translocator proteins that upon host cell contact form a pore in the host cell membrane serving as a regulated entry gate for the bacterial effectors (Figure 3A) [1–3, 8, 11–15].

Assembly of the injectisome starts with formation of the basal body by the bacterial Sec system and is followed by export of the needle proteins as early T3SS substrates. The tube-like needle then allows passage of the tip complex and translocon proteins, which are intermediate substrates. Bacterial effector proteins, representing the late T3SS substrates, are thought to be translocated into host cells through a conduit formed by needle complex and translocon.

Although electron microscopy, crystallography and biophysical techniques have provided a high resolution picture of the assembly and architecture of needle complex and sorting platform [6], major properties of the translocon such as its composition, exact localization - i.e. attached to or separated from the needle tip - or regulation have remained elusive or controversial [16].

All investigated T3SSs express two hydrophobic translocators, a major translocator harboring two and a minor translocator harboring one transmembrane domain [14, 17], Numerous studies suggested that both translocators are required for a functional translocon and assemble into a heteromultimeric pore complex [14, 17], Although T3SS translocation pores have not been visualized directly in situ, the inner opening of the pore is thought to have a diameter of 2-4 nm, as estimated by in vitro reconstitution, osmoprotection and dextran release assays [17–24]. Reconstitution in liposomes suggested that the *P. aeruginosa* translocators PopB and PopD, which are highly homologous to *Yersinia* YopB and YopD, form a hexadecameric 8:8 complex [25]. A pore complex transferred by *Y. enterocolitica* into erythrocyte membranes displayed a molecular weight of 500-700 kD [26]. Of note, considering the approximate molecular weights of 44 kD for YopB and 34 kD for YopD, a 8:8 complex of YopB and YopD would have an expected molecular weight of around 624 kD.

A previous report challenged the paradigm that effector translocation by *Yersinia pseudotuberculosis* proceeds through a continuous conduit from the bacterium directly into the target cell. Rather, it was proposed that isolated effectors located on the surface of the bacteria can translocate into target cells with the help of a separate and even an unrelated *Salmonella* T3SS [16].

It has become clear that effector translocation as well as isolated pore activities of T3SSs, such as disruption of cells and increased permeability of cell membranes, are regulated by diverse host factors including actin, Rho proteins, cholesterol, sphingolipids, coatomers, clathrin and exocyst [27–30]. In *Yersinia*, filamentous (f)-actin disruption and Rho GTPase inhibitors block translocation of effector Yops and YopB/D pore activity, whereas activation of Rho GTPases enhances effector Yop translocation [31–35]. In addition, in a feedback mechanism pathogenic yersiniae can control T3SS function through their own effectors, i.e. by modulating Rho protein activity in host cells [27].

In this work we performed high resolution fluorescence and immunogold electron microscopy to visualize translocons of *Y. enterocolitica* during host cell infection. Thereby we deciphered that translocons are connected to the remaining T3SS, that their overall number is changed dependent on the level of effector translocation and that they are formed in a specific host cell compartment.

## Results

### *Yersinia* translocators YopB and YopD localize on the bacterial cell surface after secretion by the T3SS

During infection with pathogenic yersiniae the translocators YopB and YopD are inserted into host cell membranes where they form a pore (also termed translocon) that acts as an entry gate for the *Yersinia* effectors [18, 21, 26]. To visualize the *Yersinia* translocators YopB and YopD we produced specific rabbit and rat polyclonal antibodies (S1 Fig A and B for antibody specificity). In lysates of *Y. enterocolitica* WA-314 (wild type; table 1) grown at 27°C, YopB and YopD could not be detected by immunoblot.

**Table 1.** Yersinia strains

| Strain | Relevant characteristic | Source/References |
| --- | --- | --- |
| ***Yersinia enterocolitica*** |  |  |
| **WA-314** | Wild-type strain carrying virulence plasmid pYV; serogroup O8; kanamycin resistance cassette in non-coding region of pYV-O8 | [40, 41] |
| **WA-C** | pYV-cured derivative of WA-314 | [40] |
| **WA-314ΔYopE** | WA-C harboring pYVΔYopE; Kan<sup>R</sup> | [42] |
| **WA-314ΔYopD** | WA-C harboring pYVΔYopD; Kan<sup>R</sup> | this work |
| **WA-314ΔYopB** | WA-C harboring pYVΔYopB; Kan<sup>R</sup> | this work |
| **E40 GFP-YscD** | E40 pAD4306; serogroup O9; pYVe40 (*yopO*<sub>Δ2-427</sub> *yopE*<sub>21</sub> *yopH*<sub>Δ1-352</sub> *yopM*<sub>23</sub> *yopP*<sub>23</sub> *yopT*<sub>135</sub> *egfp-yscD*); *asd*<sub>Δ292-610</sub> (auxotrophic for diaminopimelic acid) | [1] |

However, YopB and YopD proteins became detectable in bacteria grown at 37°C in high Ca^2+^ medium (non-secretion condition) and their levels further increased in bacteria grown in low Ca^2+^ medium (secretion condition; Fig 1A). It is well accepted that *Yersinia* T3SS gene expression is switched on at 37°C and that depletion of Ca^2+^ from the *Yersinia* growth medium triggers Yop secretion and boosts Yop production in a process named low Ca^2+^ response [36–38]. YopB and YopD were immunofluorescence-stained in *Yersinia* wild type at secretion condition using a procedure not supposed to permeabilize the bacterial double membrane. By confocal microscopy intense YopB and YopD fluorescence signals were seen along the bacterial circumference in essentially 100% of the bacteria (Fig 1B, C and S1 Fig C). In comparison, essentially no YopB/YopD immunofluorescence signals were found in bacteria at non-secretion condition and in bacteria at secretion condition which were treated with proteinase K (PK) before staining (Fig 1B and S1 Fig C upper panel and 1C). When immunostaining was performed in the presence of 2% sodium dodecyl sulfate (SDS) to permeabilize the bacterial double membrane, essentially all bacteria at non-secretion and secretion conditions displayed YopB and YopD signals (Fig 1B and C; S1 Fig C). In this case the YopB/YopD signals filled the whole bacterial cell rather than just the cell periphery, indicating that also intrabacterial pools of the proteins were stained. Accordingly, addition of proteinase K before staining of the bacteria under permeabilizing condition did not alter the YopB signal (Fig 1B and S1 Fig C). Total YopB signals were considerably higher in bacteria at secretion than at non-secretion condition, consistent with the immunoblot data (Fig 1A and B; S1 Fig A and C).

**Figure 1.**
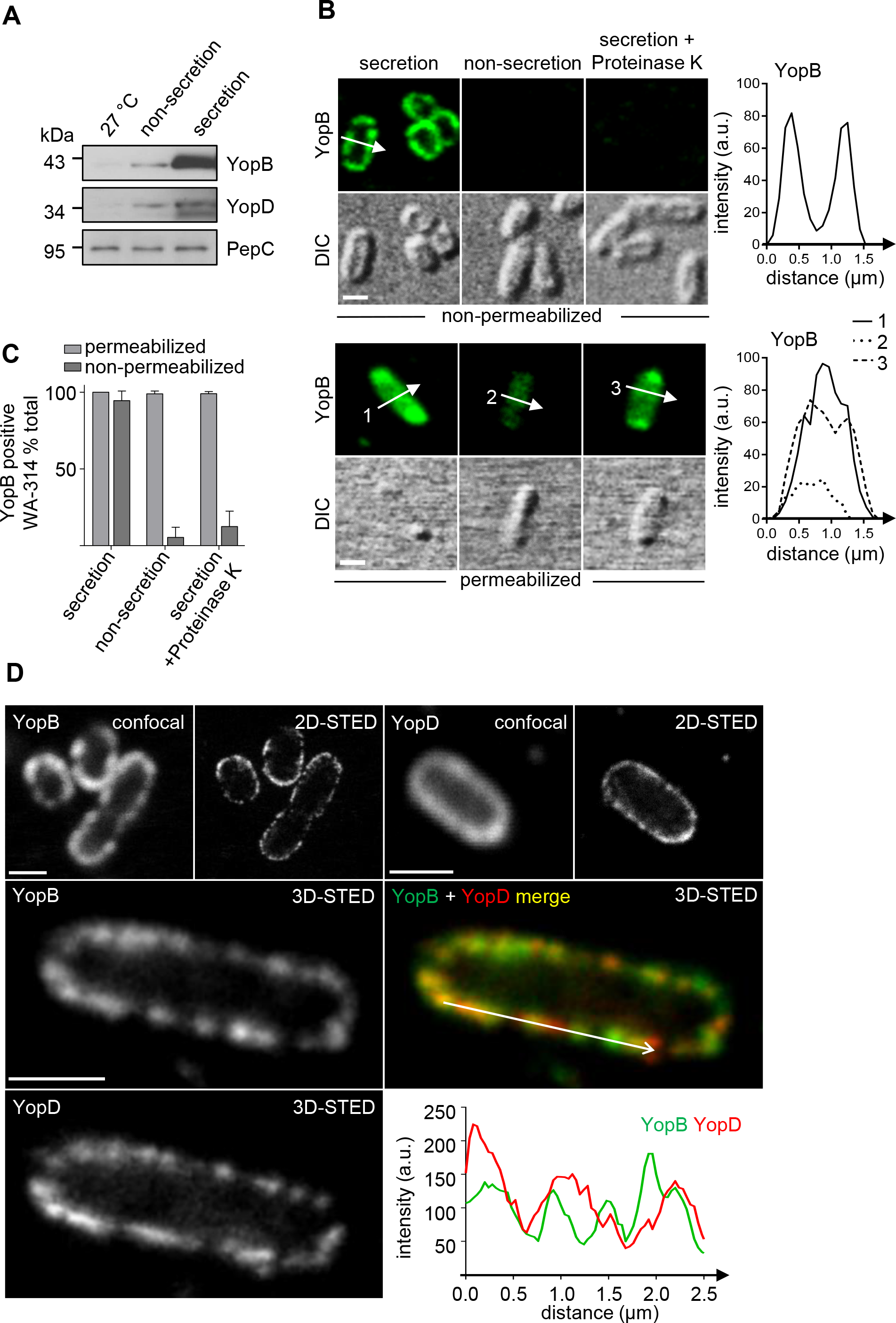
Translocators YopB and YopD decorate the *Yersinia* cell surface after secretion by the T3SS. (A) Analysis of YopB- and YopD expression. Lysates of *Yersinia* WA-314 (wild type) grown at 27°C, 37°C (high Ca^2+^/non-secretion) or 37°C (low Ca^2+^/secretion) were subjected to SDS-PAGE and analyzed by Western blot for expression of YopB, YopD or PepC, as loading control. **(B) and (C) Immunofluorescence staining of YopB in bacterial cells under secretion and non-secretion and cell permeabilizing and non-permeabilizing conditions.** Confocal immunofluorescence and corresponding differential interference contrast (DIC) images of *Yersinia* WA-314 subjected to indicated conditions. Diagrams depict fluorescence intensity profiles (arbitray units, a.u.) along the arrows in the images. Scale bars: 1 μm. **(C)** Quantitative analysis of YopB positive bacteria treated as in (B). Bars represent mean ± S.D. of n=260-800 bacteria from 2-3 independent experiments. **(D) Confocal and STED imaging of YopB and YopD.** Representative confocal and STED images (2D- or 3D-STED as indicated) of surface localized YopB and YopD in wild type *Yersinia* at secretion condition. Merge of YopB (green) and YopD (red) staining and superimposition of YopB and YopD fluorescence intensity profiles along the arrow on the bacterial surface. Scale bars: 1 μm.

To visualize YopB and YopD on the bacterial surface with higher spatial resolution, we employed stimulated emission depletion (STED) microscopy [39]. STED microscopy increases resolution of fluorescence signals to approximately 30-80 nm under the condition used (lateral resolution at 100% 2-D STED). The YopB and YopD fluorescence signals recorded with STED microscopy appeared sharper and confined to a narrow band encompassing the bacterial periphery when compared to the signals obtained with confocal microscopy (Fig 1D). Co-immunostaining revealed no systematic colocalization of YopB and YopD on the bacterial surface as demonstrated by merge of YopB and YopD 3D-STED images and by superimposed intensity plots of the YopB and YopD signals on the bacterial surface (Fig 1D).

We conclude that during the low Ca^2+^-response the secreted translocators YopB and YopD localize on the *Yersinia* cell surface but do not display a specific (co)localization pattern when investigated with high resolution fluorescence microscopy.

### YopB and YopD associate with injectisomes during cell infection

In order to visualize operational translocons, YopB and YopD were immunostained during *Y. enterocolitica* infection of HeLa cells and primary human macrophages. Confocal micrographs revealed that YopB and YopD concentrate in distinct patches in host cell associated wild type bacteria (Fig 2A and S2 Fig A). This clustered appearance as imaged by confocal microscopy was in clear contrast to the more uniform YopB/YopD distribution in secreting bacteria (Fig 1B and D). Time course experiments demonstrated that the fraction of wild type bacteria that stained positive for YopB in HeLa cells reached only about 2% at 60 min post infection (Fig 2B). By comparison, in human macrophages, which represent more physiological target cells of pathogenic yersiniae [33, 43, 44], the fraction of YopB positive bacteria amounted to around 25% at 20 min post infection (Fig 2B).

**Figure 2.**
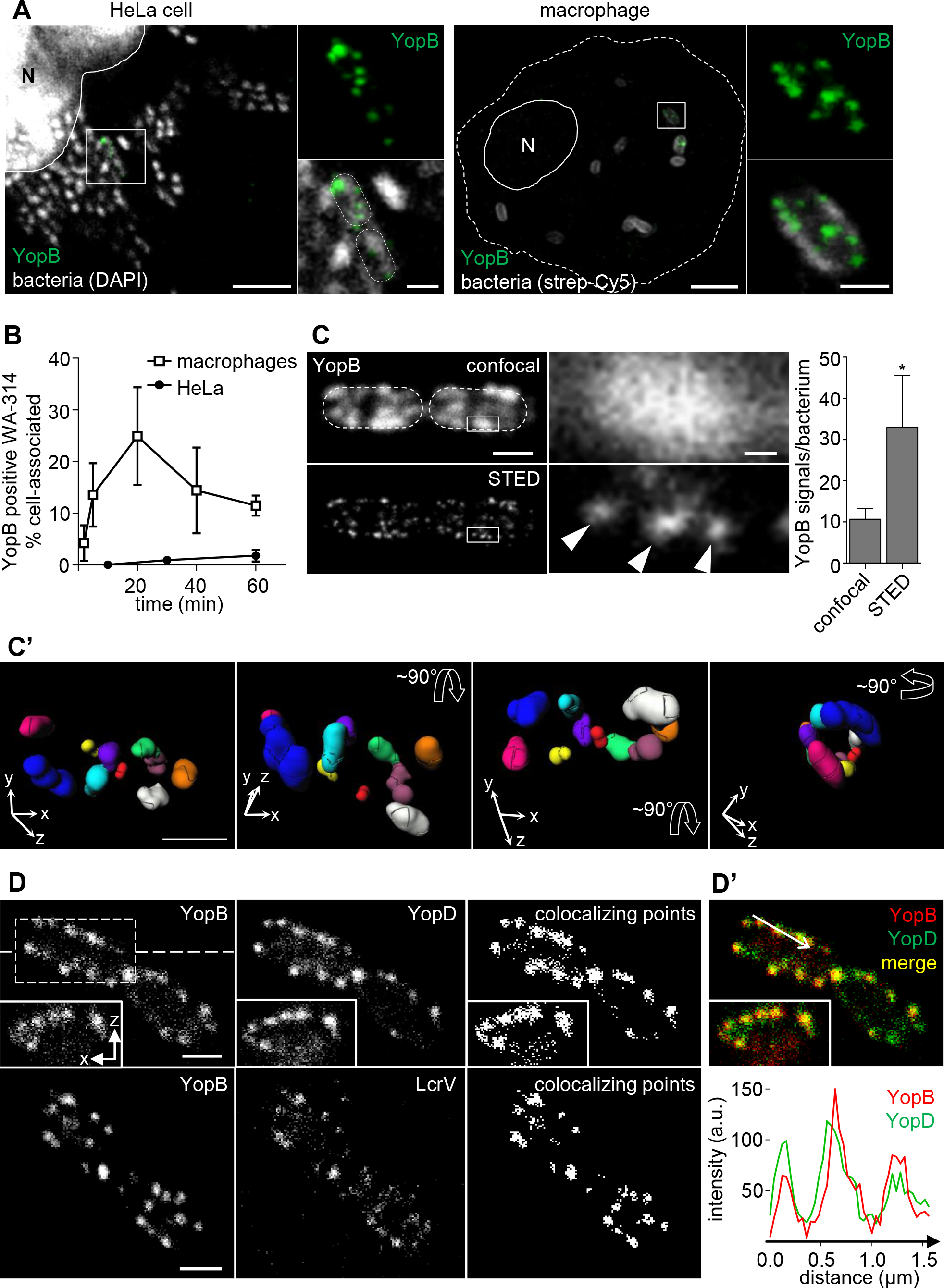
STED imaging of YopB and YopD during *Yersinia* infection of host cells. **(A) YopB concentrates in distinct patches on host cell associated bacteria.** HeLa cells (left panel) were infected with WA-314 at a MOI of 100 for 2 h. Primary human macrophages (right panel) were infected with biotinylated WA-314 at a MOI of 10 for 20 min. Cells were immunostained for YopB and either DAPI (left panel) or streptavidin-Cy5 (strep-Cy5; right panel) stained. White outlines indicate position of the nucleus (N) and dashed lines indicate host or bacterial cell circumference. Boxed regions in overviews are depicted as 2.5-fold (left) or 5-fold (right) enlargements. Scale bars: 5 μm (overviews) and 1 μm (enlargements). **(B) The fraction of YopB positive, cell-associated bacteria is dependent on the target cell type.** HeLa cells or primary human macrophages were infected with WA-314 at a MOI of 50 or 10, respectively, for the indicated time points and stained as in (A). Data represent mean ± S.D. of n=4210 bacteria associated with HeLa cells from 2 experiments or n=3047 bacteria associated with macrophages from 3 different donors **(C) YopB patches by confocal microscopy can be resolved into distinct spots by STED microscopy.** HeLa cells were infected with WA-314ΔYopE at a MOI of 100 and stained for YopB using AbberiorStarRed secondary antibody. Single z-planes of parallel confocal (upper panel) and 2D-STED (lower panel) recordings of two cell-associated bacteria are depicted in overviews. Dashed lines indicate bacterial circumference. Scale bar: 1 μm. Boxed region in overviews are depicted as 10-fold enlargements on the right. Scale bar: 0.1 μm. The number of distinct YopB signals per bacterium (WA-314YopΔE) was quantified by parallel recordings of z-stacks in confocal and 3D-STED mode. Bars represent mean ± S.D. of n=27 YopB positive bacteria from two independent experiments; *p < 0.0001 **(C’) YopB spots are organised in clusters on the bacterial surface.** Myc-Rac1Q61L transfected HeLa cells were infected with WA-314 at a MOI of 50 for 1 h and stained with anti-YopB antibody and Abberior635P secondary antibody. Z-stacks were recorded in 3D-STED mode and YopB spots on individual bacteria were subjected to image analysis (Methods). Identified clusters (consisting of at least 2 YopB spots) are projected in different colors and angles in a superresolution 3D image of a representative bacterium. Xyz-coordinate axes and arrows indicate position and horizontal or vertical rotation of the bacterium (see also S1 movie) Scale bar: 1 μm **(D) YopB/YopD and YopB/LcrV colocalize in STED images.** WA-314ΔYopE infected HeLa cells were coimmunostained for YopB (secondary antibody AlexaFluor-594) and either YopD (Abberior-StarRed) or LcrV (Abberior-StarRed). All images show representative single planes of z-stacks recorded in 3D-STED mode. Insets represent an xz projection at the level of the dashed line. Representation of colocalizing points was generated using the “Colocalization” plugin in ImageJ. Scale bar: 1 μm **(D’)** Merge (yellow) of green (YopD) and red (YopB) fluorescence. Superimposed intensity profiles of YopB and YopD fluorescence along the arrow in merge.

We next investigated the YopB signals in HeLa cells using confocal and STED microscopy in parallel. In strain WA-314ΔYopE (table 1), which was well suited for this analysis because it displayed a much higher percentage of YopB positive bacteria than wild type (see below), single patches in the confocal recordings could regularly be resolved into 3 distinct spots with the STED technology (Fig 2C). In the mean around 11 YopB positive patches and 33 YopB positive spots were detected per bacterium when investigated with confocal- and STED microscopy, respectively (Fig 2C). For a more comprehensive evaluation of the organisation of the YopB spots on the bacterial surface they were recorded in 3D-STED mode and subjected to image analysis. Spot detection and segmentation as well as cluster analysis revealed that the YopB spots are regularly organized in clusters with on average 2.8 ± 1 spots per cluster. The mean distance between the spots in one cluster was 127 ± 42 nm and the mean distance between individual clusters was 713 ± 121 nm (mean ± S.D., n=64; Methods). To exemplarily visualize the distribution of all identified clusters on a single bacterial cell, superresolution 3D images were prepared in which each segmented cluster is represented by a different color and the bacterium is viewed from different angles (Fig. 2C’ and S2 Fig B and S1 movie; Methods).

We reasoned that if the YopB and YopD spots detected by STED microscopy reflect translocons associated with the injectisome, YopB and YopD should colocalize with each other as well as with the tip complex protein LcrV, which is essential for linking YopB/YopD to the needle tip [45, 46]. YopB and YopD showed a complete colocalization in host cell associated bacteria, which was best documented when colocalizing points or intensity plots of YopB and YopD fluorescence on the bacterial surface were determined (Fig 2D and 2D’). Furthermore, STED images of YopB and LcrV co-immunostaining indicated that essentially all YopB signals are associated with LcrV signals (Fig 2D). The LcrV staining was less bright than the YopB staining which may result from the lower number of LcrV molecules per injectisome [47–49].

To further support the notion that the YopB/YopD fluorescence signals in the cell associated bacteria represent translocons attached to needle tips, we investigated the spatial coupling of YopB/D to the basal body component YscD at high resolution. Considering the published data on *Yersinia* basal body, needle and tip complex length as well recent work in *Salmonella*, the N-terminal domain of YscD - or of its homologues - is located in approximately 100 nm distance from the needle tip (Fig 3A, [13, 50–53]). For this experiment HeLa cells were infected with *Y. enterocolitica* strain E40 GFP-YscD [1] (table1) expressing a N-terminal EGFP-fusion of YscD and immunostained for YopB or YopD. Because GFP-YscD did not produce a high enough fluorescence signal for STED microscopy, the alternative high resolution microscopy technique structured illumination microscopy (3D-SIM, resolution to ~100 nm in x-y plane, [54, 55]) was employed to visualize GFP-YscD and YopB/D in parallel. SIM revealed single YopB dots that could clearly be separated from each other and from neighboring patches of GFP-YscD (Fig 3B). A fluorescence intensity plot of a representative YopB/GFP-YscD pair indicated a distance of 90 nm between the fluorescence maxima (Fig 3B). A more comprehensive nearest neighbour analysis indicated distances between the GFP-YscD patches and YopB/D dots of around 110 nm (109 ±4 nm; n=424 pairs evaluated; Methods; Fig 3C).

**Figure 3.**
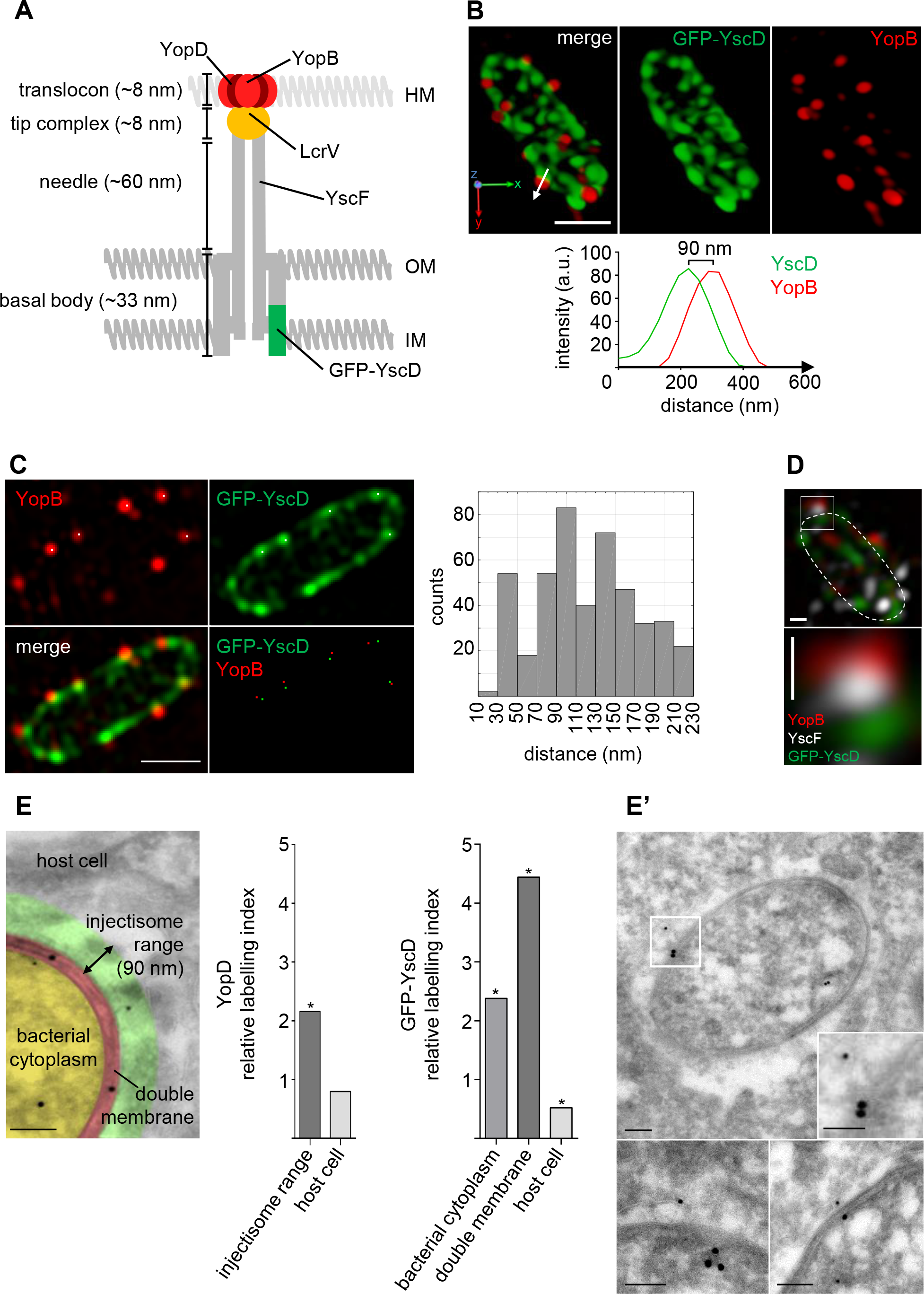
Spatial coupling of translocators YopB/YopD to basal body component YscD. **(A) Schematic representation of the *Yersinia* type 3 secretion injectisome.** Positions of YopB/YopD, LcrV, YscF and GFP-YscD and the approximate length of basal body, needle, tip complex and translocon of a virtual injectisome are indicated [13, 52, 53, 69]. Host membrane (HM), outer membrane (OM), inner membrane (IM). **(B) SIM separates YopB dots from near neighboring patches of GFP-YscD.** HeLa cells were infected with *Y. enterocolitica* E40 GFP-YscD, stained with anti-YopB antibody and z-stacks of YopB positive bacteria were recorded with SIM. A 3D reconstruction of a representative bacterium is depicted (see also S2 movie). Scale bar: 1 μm. Fluorescence intensity profiles along the longitudinal axis (arrow) of a GFP-YscD/YopB pair indicate a distance of 90 nm between fluorescence maxima. **(C) Nearest neighbor distance analysis of YopB/D and GFP-YscD fluorescence signals.** HeLa cells were infected with *Y. enterocolitica* E40 GFP-YscD, stained with anti-YopD or anti-YopB antibody and z-stacks of YopB/D positive bacteria were recorded with SIM. Coordinates of clearly discernible individual YopB or YopD and GFP-YscD signals were retrieved from single z-planes using ImageJ plugin TrackMate. Distances of individual YopB or YopD dots to the nearest GFP-YscD patches were analyzed by a nearest neighborhood analysis in Matlab and are depicted as histogram (mean ± S.D. was 109 ± 4 nm; 424 pairs measured; Methods). Corresponding YopB/YscD pairs (red and green) complying with a distance limit of 200 nm were plotted back on the original fluorescence images (white dots in GFP-YscD and YopB images) for illustration. Scale bar: 1 μm. **(D)** Double immunostaining of YscF and YopB in E40 GFP-YscD infected HeLa cells followed by SIM reveals conjugated fluorescence triplets displaying basal body (green), needle (white) and translocon (red). Dashed line indicates bacterial circumference. Scale bars: 100 nm. **(E) Enrichment of YopD within the injectisome range.** Transmission electron microscopy (TEM) images were compartmentalized into bacterial cytoplasm (yellow), bacterial double membrane (red), injectisome range (green, 90 nm from bacterial outer membrane) and adjacent host cell (grey). Scale bar: 100 nm. Relative labelling indices of YopD were determined for the injectisome range and the adjacent HeLa cell (n=80 golds evaluated). Relative labelling indices of GFP-YscD were calculated for bacterial cytoplasm, bacterial double membrane and the host cell (n=580 golds evaluated). *p < 0.00001 different from expected distribution (S1 Tables A-C, analysis according to [75]) **(E’) Double immunogold staining of YopD and GFP-YscD.** HeLa cells expressing myc-Rac1Q61L were infected with *Y. enterocolitica* E40 GFP-YscD and subjected to ultrathin sectioning and immunogold staining of GFP-YscD (15 nm particles) and YopD (10 nm particles). TEM images show exemplary membrane engulfed bacteria with a GFP-YscD/YopD configuration (≤ 140 nm apart; Methods) indicative of a translocon containing/fully assembled injectisome. Scale bars: 100 nm.

Having successfully resolved fluorescence signals from translocon and basal body proteins that are located in distant parts of the same injectisomes, we next coimmunostained the needle protein YscF with YopB in strain E40 GFP-YscD. YscF forms the needle that connects translocon and basal body in injectisomes (Fig 3A). SIM revealed tripartite complexes made up of YopB, GFP-YscD and YscF, whereby the fluorescence signal for YscF was sandwiched between the YopB and YscD signals (Fig 3D). Thus, by high resolution SIM we were able to resolve three proteins located in different parts of *Yersinia* injectisomes.

We next aimed to visualize YopD and GFP-YscD embedded in their native bacterial and host cellular environment. Transmission electron microscopy (TEM) of ultrathin sections combined with cryoimmunolabelling allows high resolution localization of proteins within their cellular context [56, 57]. Electron-dense protein-A/gold particles of different sizes (usually 5-15 nm) additionally permit to localize different proteins in the same sections [58]. We infected Rac1Q61L expressing HeLa cells, in which the expression of *Yersinia* injectisomes is strongly enhanced (see below), with strain E40 GFP-YscD and immunolabelled YopD and GFP-YscD with 10 nm and 15 nm gold particles, respectively. The labelling density of the 10 nm gold particles marking YopD was much higher in the bacterial cell (cytoplasm and double membrane; relative labelling index (RLI: 2.73) than in the extrabacterial area (RLI: 0.42; Fig 3E and S1 table A). Thus, the observed distribution of the 10 nm gold particles reflects the expected YopD localization in section staining, which includes the injectisome and the intrabacterial pools (S1 Fig C). We next aimed to assess if the extrabacterially located 10 nm gold particles could represent YopD in translocons at the tip of injectisome needles. We assumed that in this case the gold particles should be located within a range of 90 nm from the bacterial outer membrane which we defined as injectisome range, reflecting the cumulative dimensions of the injectisome needle (~60 nm), tip complex (~8 nm) and antibody/protein A complex used for immunogold staining (~20 nm; Fig 3E). Distances less than 90 nm may occur depending on the geometry and spatial orientation of the type III secretion machines in the 2D analysis.

YopD labelling within the injectisome range was significantly enriched (RLI: 2.16) compared to the remaining host cell area (RLI: 0.79; Fig 3E and S1 table B) suggesting that it represents translocons associated with injectisomes.

The labelling density of the gold particles marking GFP-YscD was highest in the bacterial double membrane (RLI: 4.44) followed by the bacterial cytoplasm (RLI: 2.38) and was only minor in the extrabacterial space (RLI: 0.52; Fig 3E and S1 table C). This distribution reflects the expected localization of GFP-YscD in assembled basal bodies in the inner bacterial membrane and in an intrabacterial YscD pool. We finally employed co-immunogold staining to detect GFP-YscD and YopD localized within the same injectisomes. Considering the dimensions of the injectisome (length of ~100 nm; Fig 3A) and the two antibody/protein A complexes (length of 2 × 20 nm = 40 nm), we assumed that 10 nm gold particles (YopD label) that lie extrabacterially and in an at most ~140 nm distance from 15 nm gold particles, that themselves are located in the bacterial double membrane (YscD label), belong to the same fully assembled injectisome.

Consistent with this notion we repeatedly identified configurations of 10 nm and 15 nm gold particles fulfilling these premises (Fig 3E’). Of note, the 10 nm gold particles were often found in close proximity to the host cell membrane surrounding the bacteria (Fig 3E’). Thus, the observed distributions of the 10 nm and 15 nm gold particles in bacteria surrounded by host cell membranes reflect the expected locations of YopD and GFP-YscD at the tip and basal body, respectively, of injectisomes.

In summary, we conclude that the *Yersinia* translocons can be visualized in fully assembled injectisomes by superresolution fluorescence microscopy (SIM and STED) and immunogold TEM techniques, the latter method also suggesting that translocons are located in a specific cellular compartment

### Rac1 activity and host cell type regulate *Yersinia* translocon formation

The visualization of putatively operable translocons in host cells prompted us to test how translocon number or distribution correlate with the level of effector translocation. To enhance Yop effector translocation into HeLa cells, cells were infected with the YopE-deficient strain WA-314ΔYopE or cells expressing the constitutively active Rho GTP-binding protein Rac1Q61L were infected with wild type *Yersinia.* Under these conditions Yop-translocation rates increase 5-10-fold [35] which is a consequence of the elevated Rac activity in the host cells [35, 59, 60]. In the case of WA-314ΔYopE the elevated Rac activity is caused by the diminished Rac inhibition that is normally imposed by the Rho GTPase-activatingprotein YopE [61, 62], Notably, the fraction of YopB positive bacteria increased from around 2% in wild type infected cells to 30% in WA-314ΔYopE infected cells and also reached around 30% in the Rac1Q61L expressing and wild type infected cells at 1 h post infection (Fig 4A).

**Figure 4.**
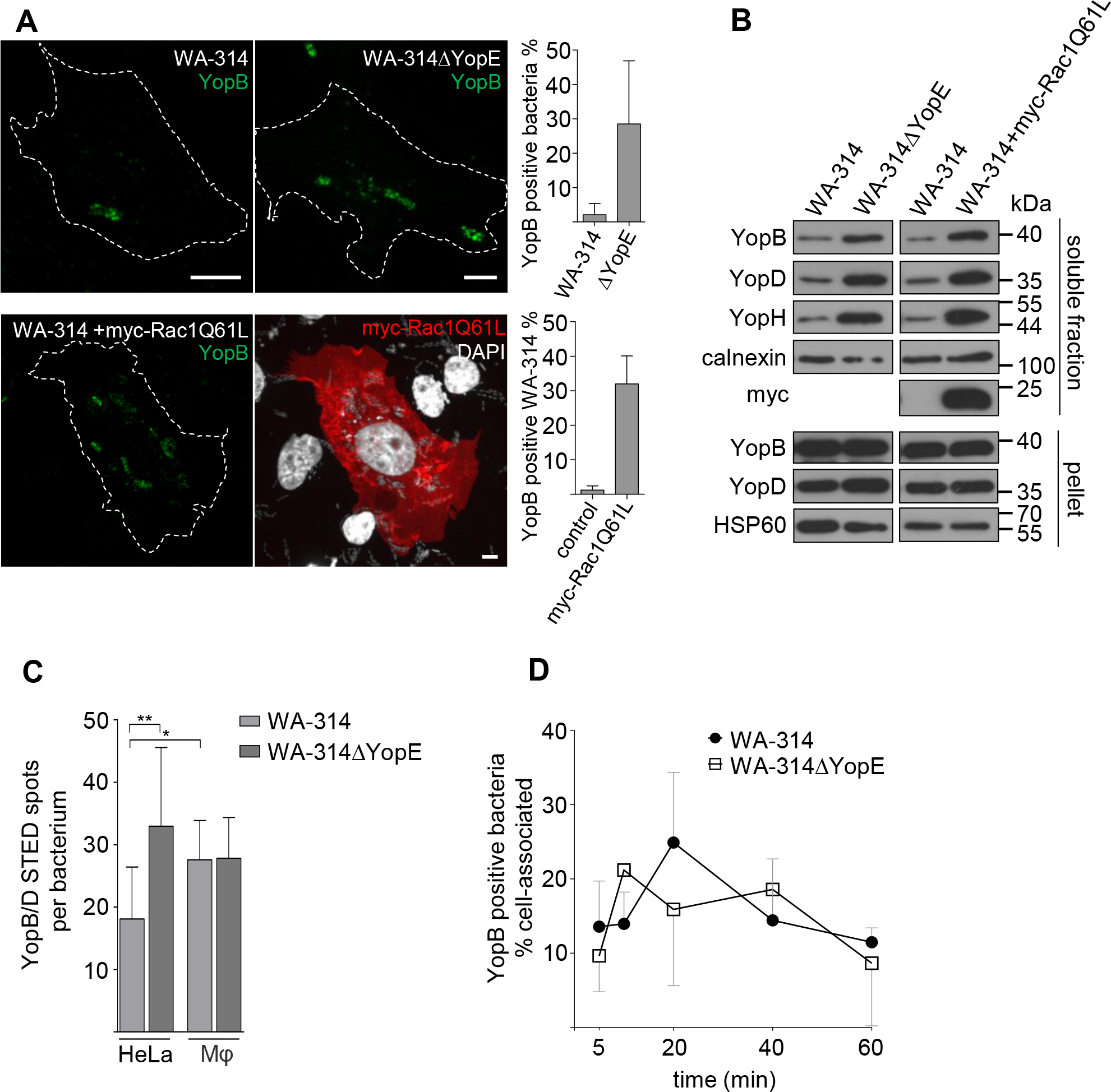
Racl activity and host cell type regulate translocon formation. **(A) Lack of Racl inhibitor YopE and Rac1Q61L overexpression enhance translocon formation.** Native HeLa cells were infected with WA-314 or WA-314ΔYopE and myc-Rac1Q61L overexpressing HeLa cells were infected with WA-314 at a MOI of 50 for 1 h. Cells were immunostained for YopB and myc as well as DAPI stained. Dashed lines indicate host cell circumference. Scale bars: 5 μm. The percentage of YopB positive bacteria of total cell associated DAPI-stained bacteria was determined. Bars represent mean ± S.D. of YopB positive bacteria per cell-associated bacteria. In total n=6505 WA-314 and n=5640 WA-314ΔYopE bacteria were investigated in 4 independent experiments (upper graph); n=3423 bacteria (WA-314) associated to control transfected cells and n=7882 bacteria (WA-314) on myc-Rac1Q61L transfected cells were investigated in 3 independent experiments (lower graph). **(B) Digitonin extraction reveals increased incorporation of YopB and YopD into host cell membranes upon Rac1Q61L overexpression or lack of Racl inhibitor YopE.** Experimental conditions as in (A) but with a MOI of 100. HeLa cells were lyzed with digitonin and resulting supernatants (containing membrane integrated and soluble Yops from host cells) and cell pellets (containing intact bacteria and digitonin insoluble cell components) were analyzed with Western blot for the indicated proteins. YopH serves as marker for effector translocation, myc indicates myc-RaclQL61 expression, calnexin serves as host cell loading control and HSP60 serves as bacterial loading and lysis control. Data are representative of 3 independent experiments. **(C) Quantification of YopB spots per bacterium using STED microscopy.** Experimental conditions for Hela cell infection as in (A). Human primary macrophages (Mφ) were infected with WA-314 or WA-314ΔYopE at a MOI of 20 for 20 min. Cells were immunostained with anti-YopB antibody followed by AbberiorStarRed or Abberior635P secondary antibody. Z-stacks of YopB positive bacteria were recorded in 3D-STED mode. The number of YopB spots per bacterium was analyzed using Imaris software (Methods). Bars represent mean ± S.D. of n=20 WA-314 (Hela), n=27 WA-314ΔYopE (Hela), n=77 WA-314 (Mφ) and n=7 WA-314AYopE (Mφ) bacteria investigated in 2-3 independent experiments. *p < 0.05, **p < 0.01 **(D) YopB positive WA-314 and WA-314ΔYopE in macrophages at different infection times.** Primary human macrophages were infected with WA-314 or WA-314AYopE at a MOI of 10 for the indicated time periods. The percentage of YopB positive bacteria of total cell associated bacteria was calculated. Data represent mean ± S.D. of n=3047 WA-314 and n=1822 WA-314ΔYopE in 2-3 independent experiments.

As an alternative method to detect host membrane inserted translocons we employed a digitonin-based release assay. Digitonin extracts host cell associated Yops but not intrabacterial Yops and has hitherto been used to assay effector-Yop translocation [63]. Consistent with an increased deposition of translocons in the infected cells, the amounts of YopB and YopD extracted by digitonin were increased in the WA-314ΔYopE infected cells as well as in the wild type infected and Rac1Q61L expressing cells when compared to the respective controls (Fig 4B). The extracted amounts of YopB and YopD correlated with the extracted amount of YopH, the latter serving as a measure for effector-Yop translocation (Fig 4B). In comparison, the levels of YopB, YopD and HSP60 in the pellet fraction (representing the bacterial protein pool) were similar in all conditions.

Thus, in infected HeLa cells stimulation of Yop translocation is associated with a large increase in the number of bacteria forming translocons which causes enhanced translocon incorporation into cell membranes.

We next tested whether the number of translocons per bacterium, as identified by YopB fluorescence spots in STED images, was also altered under conditions of increased Yop translocation. In strain WA-314ΔYopE the number of translocons was nearly twice as high as that in wild type upon infection of HeLa cells (Fig 4C). Furthermore, in macrophages wild type bacteria showed a significantly higher number of translocons than in HeLa cells reaching similar values as WA-314ΔYopE in HeLa cells (Fig 4C). There was no difference in the number of translocons formed between wild type and WA-314ΔYopE in macrophages (Fig 4C) excluding that the WA-314ΔYopE strain intrinsically produces more translocons. Interestingly, in macrophages there was also no difference in the fraction of translocon positive bacteria between wild type and WA-314ΔYopE (Fig 4D).

These results indicate that host factors not only regulate the number of bacteria producing translocons and thereby the level of translocons within cell membranes but also can affect the number of translocons/injectisomes per bacterium. They also indicate that macrophages can trigger pathogenic yersiniae much more effectively than HeLa cells to produce translocons. Of note, upon activation of Rac HeLa cells acquire the same potency as macrophages in stimulating translocon formation whereas translocon formation in the already highly effective macrophages appears not to be affected (Fig 2B and 4D).

### Formation of translocons is triggered in a prevacuole, a PI(4,5)P2 enriched host cell compartment

The TEM images showed translocons in yersiniae enclosed by host cell membranes. This prompted us to test whether formation of translocons is triggered in a specific host cell compartment Previously it was described that during cell invasion avirulent yersiniae enter a precompartment in which they are accessible to externally administered small proteins or compounds (MW 50 kD) but not to antibodies (MW around 150 kD, [64, 65]). This compartment was named a prevacuole and shown to be enriched in the phospholipid PI(4,5)P2 [64],

When Hela cells were infected with biotinylated wild type *Y. enterocolitica* for 60 min and bacterial accessibility to externally administered streptavidin (MW 53 kD) or *Yersinia* specific antibodies was assessed by fluorescence staining, three different staining patterns were found. The bacteria were either accessible to antibodies and streptavidin (denominated outside), inaccessible to antibodies but accessible to streptavidin (intermediary) or inaccessible to both, antibodies and streptavidin (inside; Fig 5A and 5A’). Quantitative analysis showed that about 70% of the YopB positive bacteria were located in the intermediary compartment and around 25% in the inside compartment but only a negligible fraction (less than 5%) was present in the outside localization at 1 h post infection (Fig 5A”).

**Figure 5.**
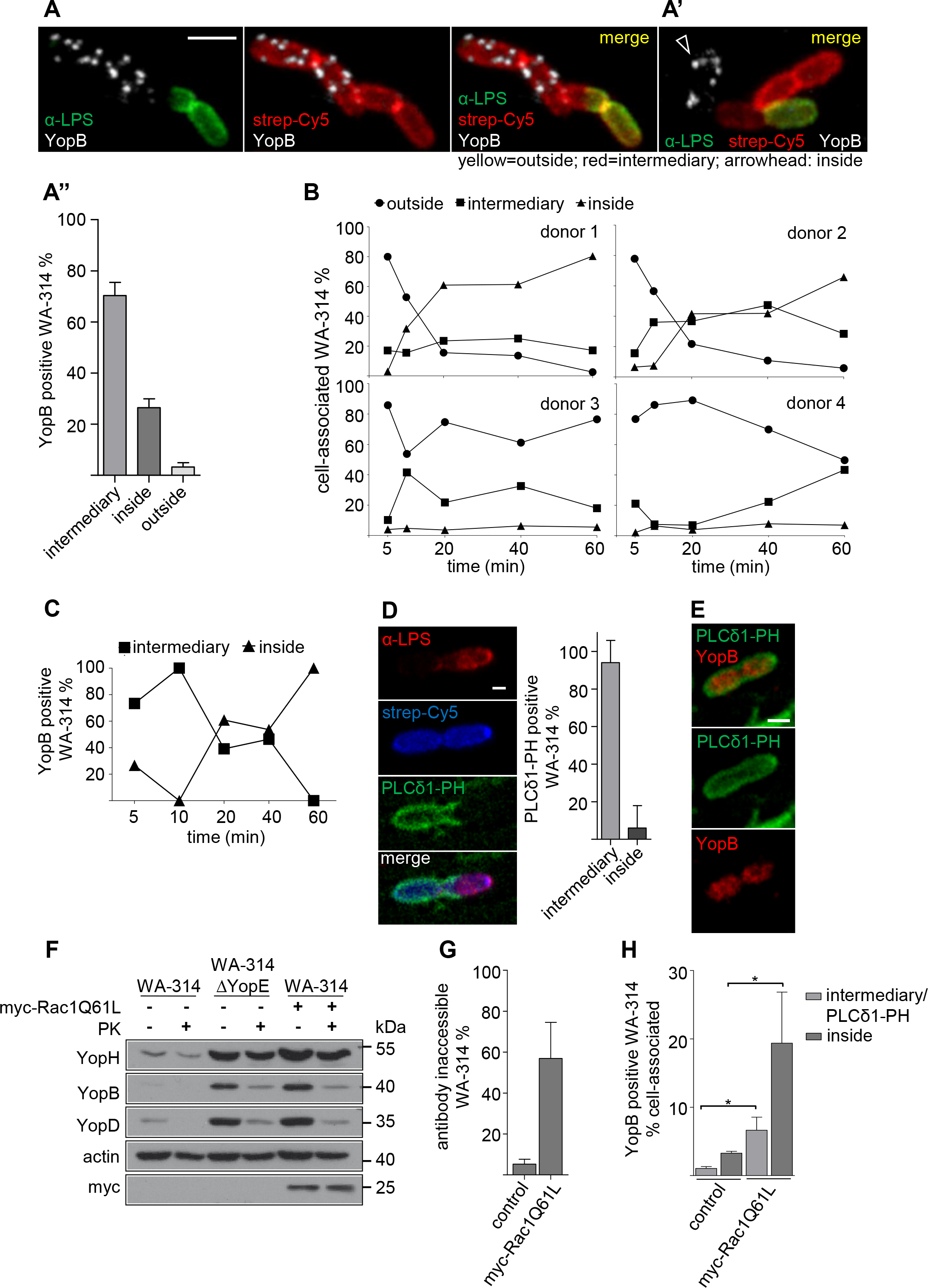
Formation of translocons is triggered in a prevacuole, a PI(4,5)P2 enriched intermediary host cell compartment. (A-A”) YopB positive bacteria reside in an intermediary compartment. Hela cells were infected with surface-biotinylated WA-314 at a MOI of 20 for 60 min and were immunostained using anti-LPS antibody and stained with streptavidin-Cy5 without cell permeabilization. Then cells were permeabilized and immunostained with anti-YopB antibody. **(A)** Images of a bacterial chain showing YopB staining only in the part of the chain located in the intermediary compartment (red in merge) but not in the outside location (yellow in merge). **(A’)** Image of a YopB positive bacterium (white) in the inside compartment (no red or green). Scale bar: 1 μm **(A”)** Distribution of YopB positive bacteria between intermediary, inside and outside compartments. Bars represent mean ± S.D. of n=95 YopB positive bacteria recorded in 2 independent experiments. **(B) Bacterial passage from the outside into the intermediary and inside compartments of macrophages.** Primary human macrophages of donors 1-4 were infected with biotinylated WA-314 at a MOI of 10 for the indicated time points and stained with anti-LPS antibody, streptavidin-Cy5 and DAPI to score their localization to the indicated compartments as in (A-A”). Data represent mean of at least n=1000 bacteria investigated per donor. **(C) YopB positive bacteria passage from the intermediary to the inside compartment in macrophages.** Experimental conditions as in (B) but with no LPS but YopB immunostaining. Data represent mean of n=952 YopB positive bacteria investigated in 1 donor. **(D) Intermediary compartment is marked by PLCδ-PH-GFP.** Hela cells expressing PLCδ-PH-GFP and myc-Rac1Q61L were infected with WA-314 at a MOI of 10 for 60 min, stained with anti-LPS antibody and streptavidin-Cy5. Bacteria with PLCδ-PH-GFP enrichment were judged for their localization as in (A-A”). Bars represent mean ± S.D. of n=398 PLCδ-PH-GFP positive bacteria investigated in 1 experiment **(E) PLC8-PH-GFP enrichment around YopB positive bacteria in human macrophages.** Primary human macrophages expressing PLCδ-PH-GFP were infected with WA-314 at a MOI of 20 for 20 min and immunostained for YopB. **(F) External addition of proteinase K to infected HeLa cells degrades YopB and YopD but not YopH.** Control or myc-Rac1Q61L expressing HeLa cells were infected with WA-314 or WA-314ΔYopE at a MOI of 100 for 1 h as indicated. Cells were incubated with PK for 20 min. PK was inactivated with PMSF, cells were extracted with digitonin and analyzed by Western blot for YopB and YopD, YopH (translocated effector protein), myc (myc-Rac1Q61L expression) and actin (loading control). Data are representative of 3 independent experiments. **(G) Rac1Q61L overexpression increases uptake of WA-314 into HeLa cells.** Control or myc-Rac1Q61L expressing HeLa cells were infected with WA-314 at a MOI of 20 for 1 h, fixed and stained with anti-LPS antibody (antibody accessible bacteria in outside location) and then permeabilized and stained with the same anti-LPS antibody in combination with a different secondary antibody (antibody inaccessible bacteria in intermediary and inside compartments). Bars represent mean ± S.D. of n=1609 bacteria on control transfected cells and n=3482 bacteria on myc-Rac1Q61L transfected cells investigated in 3 independent experiments. **(H) Rac1L61 expression significantly increases the number of YopB positive bacteria in the intermediary and inside compartments.** HeLa cells expressing PLCδ-PH-GFP and either empty vector (control) or myc-Rac1Q61L were infected with WA-314 at a MOI of 50 for 1 h and stained with anti-YopB antibody. The number of YopB positive bacteria as fraction of total cell associated bacteria and their distribution between the intermediary (PLCδ-PH-GFP positive) and inside (PLCδ-PH-GFP negative) compartments was determined. Bars represent mean ± S.D. of n=1820 bacteria on control transfected cells and n=6020 bacteria on myc-Rac1Q61L transfected cells investigated in 3 independent experiments.

To find out in which macrophage compartment translocons are formed, we assayed YopB fluorescence signals of bacteria during their passage into the intermediary and inside compartments. For this macrophages prepared from four different donors were infected with wild type and investigated during a 60 min time period. The bacteria started to enter the intermediary compartment before 5 min of infection (Fig 5B) and in all but one preparation of macrophages (donor 4, Fig 5B) the fraction of bacteria that localized to the intermediary compartment remained stable between 20 to 60 min post infection. At 60 min post infection 20 to 40% of the cell associated bacteria still resided in the intermediary compartments (Fig 5B). Notably, the fraction of bacteria that further progressed to the inside compartment differed widely among the different macrophage preparations. It amounted to 60-80% in macrophages from donors 1 and 2 and was about 5% in macrophages from donors 3 and 4 at 60min post infection (Fig 5B). Thus in macrophages from some individuals only a minimal fraction of wild type *Yersinia* transits from the intermediary to the inside compartment, whereas in macrophages from other donors the bacteria readily proceed to the inside compartment.

In a preparation of macrophages in which the bacteria mostly end up in the inside compartment (similar to macrophages from donor 1, Fig 5B) it was first verified that bacteria in the outside location do not show YopB staining like already seen in HeLa cells (no YopB positive outside bacteria in 50 macrophages from two experiments investigated). At 5-10 min post infection essentially all YopB positive bacteria were found in the intermediary compartment and thereafter the fraction of YopB positive bacteria decreased in the intermediary compartment and in parallel increased in the inside compartment (Fig 5C). These results clearly indicate that translocons are formed in the intermediary compartment of macrophages and then are carried on to the inside compartment

That the translocon proteins in the intermediary compartment are accessible to externally added proteins was confirmed with a modified digitonin lysis assay. Proteinase K (MW 29 kD) was added to *Yersinia* WA-314 or WA-314ΔYopE infected HeLa cells and was then neutralized with phenylmethylsulfonyl fluoride (PMSF) prior to digitonin lysis and Western blot analysis of the cells. Under these conditions YopB and YopD were largely degraded whereas the effector YopH and actin remained unchanged (Fig 5F). It was verified in control experiments that with the proteinase K amounts employed Yops can principally be degraded and that addition of PMSF abrogates proteinase K activity (S3 Fig A, Methods).

The prevacuole formed during invasion of avirulent *Y. pseudotuberculosis* into COS1 cells was characterized by accumulation of the PI(4,5)P2 sensor PLCδ-PH-GFP [64]. Accumulation of PLCδ-PH-GFP also marked the intermediary compartment in wild type infected HeLa cells (Fig 5D) and accumulated around translocon positive wild type bacteria in human macrophages (Fig 5E).

We finally hypothesized that the strong enhancement of translocon formation in HeLa cells upon Rac1 activation may be due to the capability of Rac to stimulate bacterial uptake into the intermediary prevacuolar compartment [66–68].

In fact, in HeLa cells overexpressing myc-Rac1Q61L around 60% of wild type bacteria became inaccessible to antibodies compared to around 10% in control cells after 1 h of infection (Fig 5G). This resulted in a 6-fold higher number of YopB positive bacteria both, in the intermediary and inside compartments of myc-Rac1Q61L overexpressing cells when compared to controls (Fig 5H).

Altogether we conclude from this set of experiments that the formation of *Yersinia* translocons is triggered in a PI(4,5)P2 enriched permissive cell compartment, which is protected from large extracellular proteins like antibodies. Primary human macrophages readily internalise the bacteria in this permissive compartment, which is most certainly the reason why they so effectively stimulate translocon formation. In comparison, epithelial cells like HeLa cells possess a low intrinsic activity to internalise the bacteria in the permissive compartment. However, uptake of the bacteria into the permissive compartment, translocon formation and effector translocation can be dramatically stimulated to values reached in primary macrophages by activation of the Rho GTP binding protein Rac1.

## Discussion

The two hydrophobic translocators present in most T3SSs and the pore complex/translocon that these proteins form in host cell membranes are particularly difficult to investigate. This is amongst others due to the highly elaborate transit of the translocators from the bacterial interior where they have to be in a soluble form, through the T3SS needle when they are in an unfolded state, up to their dynamic interaction with the needle tip. The tip complex is supposed to orchestrate integration of the translocators into the host cell membrane, a process that presumably is accompanied by refolding and heteromultimeric assembly of the proteins. Because it is localized at the interface of the bacterial T3SS and the target cell membrane, the translocon is unavoidably controlled by host cell factors that determine the composition and function of cell membranes.

A recent superresolution fluorescence microscopy study of *Salmonella* Typhimurium without host cell contact described clusters of the basal body protein PrgH, a *Yersinia* YscD analogon, with a width of 46 nm [50]. The 12 PrgH clusters found on average per *Salmonella* cell were shown to represent individual needle complexes whereby the mean distance between the PrgH fluorescent signal and the signal of the tip complex protein SipD, a *Yersinia* LcrV analogon, was determined to be 101 nm [50].

Cryo-Electron Tomography (Cryo-ET) analysis of *Y. enterocolitica* (mini)cells also provided an estimation of the number and organization of needle complexes expressed without host cell contact. In tomograms of single *Y. enterocolitica* cells 6.2 injectisomes were detected on average whereby it was calculated that the employed technique underestimates the total number of injectisomes by a factor of 2-3. Cryo-ET also demonstrated that separate fluorescence signals of *Yersinia* needle complexes seen in the confocal microscope contained in the mean 2.5 injectisomes organized in clusters. Within these clusters the injectisomes were about 100 nm apart whereas more randomly distributed injectisomes showed distances of about 400 nm [13]. These numbers are in good agreement with the 18-33 translocons implying fully assembled injectisomes identified by superresolution fluorescence microscopy in *Y. enterocolitica*. They also comply well with the average distance of 127 nm between individual translocons in clusters as well as with the mean distance of 716 nm measured between the clusters. Our data further suggest that the number of operable translocons on the bacteria increases in parallel to effector translocation and is significantly higher in macrophages than in epithelial cells. In fluorescence patches of translocator proteins visualized with confocal microscopy we were able to resolve on average 3 separate translocator spots by STED, corresponding well to the 2-3 injectisomes found with Cryo-ET in fluorescence clusters of injectisome components [13]. The previously reported numbers of injectisomes in secreting bacterial cells, their dimensions and distances among each other is therefore highly concordant with the respective features of the *Yersinia* translocons described here.

Until recently a direct view of fully assembled injectisomes including translocons has not been reported. A preprint of a Cryo-ET study provided a first direct view of the T3SS translocon of *Salmonella* minicells during host cell infection [69]. The EM pictures clearly indicate that the *Salmonella* translocon is connected to the T3SS needle and is in parallel embedded in the host cell membrane and protrudes towards the host cell cytoplasm. The *Salmonella* translocons are ~13.5 nm in diameter and 8 nm thick whereby the part of the translocon protruding into the target cell creates a hole which may represent the inner opening through which the effectors enter the cell [69].

In this study we employed high resolution STED and SIM fluorescence microscopy to resolve translocon, needle and basal body proteins of *Yersinia* injectisomes during cell infection and come to the conclusion that *Yersinia* injectisomes also form a continuous conduit from the bacterial to the target cell cytoplasm. However, our findings do not exclude the additional existence of translocons operating independently of the remaining injectisome as was proposed recently [16]. Although our high resolution fluorescence data combined with double immunogold staining cannot provide high fidelity structural information like the Cryo-ET technique, fluorescence techniques are accessible to a larger number of researchers and form the basis for future live imaging studies. Moreover, our experiments were not limited to minicells, which do not translocate effectors, and we therefore could identify the role of *Yersinia* effector YopE in regulating translocon formation. Similarly, because we were not restricted to investigation of thin peripheral host cell areas as with Cryo ET, we could characterize a dynamic host cellular compartment that promotes translocon formation. Our study thereby provides an explanation for the reported stimulatory effect of Rho GTP-binding proteins on effector translocation by *Yersinia* [34, 35, 59–62]. We propose that Rac1 enhances uptake of yersiniae into the prevacuole of host cells which stimulates formation of operable injectisomes and effector translocation.

## Material and Methods

### Materials

All standard laboratory chemicals and supplies were purchased from Roth (Karlsruhe, Germany), Sigma-Aldrich (Steinheim, Germany) or Merck (Hohenbrunn, Germany) unless indicated otherwise.

### Plasmids

The following plasmids were described previously: PLCδ1-PH-GFP was provided by T. Balla (National Institutes of Health, Bethesda, MD). The myc-Rac1Q61L plasmid was kindly provided by Dr. Pontus Aspenström (Uppsala University, Uppsala, Sweden) and pRK5myc was purchased from Clontech.

### Antibodies

Polyclonal rabbit anti-YopB (aa 1-168) and anti-YopD (aa 150-287) as well as rat anti-YopB (aal-168) and anti-YscF antibodies were produced by immunization of the animals with the respective purified GST-fused proteins (animal research project A10a 675). For immunofluorescence staining, sera were affinity purified by binding either to the suitable GST-fused recombinant antigens bound to glutathione beads or to antigens released by *Y. enterocolitica* WA-314 that were run on SDS-polyacrylamide gel electrophoresis (PAGE) and blotted onto polyvinylidine fluoride (PVDF) membranes (Immobilon-P, Millipore, Schwalbach, Germany). Anti-LcrV, anti-YopH and anti-PepC [70] rabbit polyclonal sera were a gift of Jürgen Heesemann (Max von Pettenkofer-Institute, Munich, Germany).

Primary antibodies and their sources were: rabbit polyclonal anti-*Y. enterocolitica* 0:8 (Sifin, Berlin, Germany); rabbit polyclonal anti-calnexin (Enzo, Lörrach, Germany); rabbit polyclonal myc (Cell Signaling, Cambridge, UK); mouse monoclonal anti-actin (Millipore, Schwalbach, Germany); mouse monoclonal anti-HSP60 and anti-streptavidin-Cy5 (Thermo Fisher Scientific, Waltham, USA); biotin conjugated goat polyclonal anti-GFP (Rockland, Limerick, USA).

Secondary anti-IgG antibodies and their sources were: Alexa488 chicken anti-rabbit and goat anti-rat, Alexa568 goat anti-rabbit and goat anti-rat, Alexa647 goat anti-rabbit, Alexa594 chicken anti-rat (Molecular Probes, Karlsruhe, Germany). AbberiorStar580 donkey anti-rabbit, AbberiorStarRed donkey anti-rabbit and goat anti-rat, AbberiorStar635P goat anti-rabbit (Abberior, Göttingen, Germany). Rabbit polyclonal anti-biotin (Rockland, Limerick, USA). Protein A gold was purchased from G. Posthuma (University Medical Center Utrecht, Netherlands). Horseradish peroxidase linked sheep anti-mouse, donkey anti-rabbit and goat anti-rat (GE Healthcare, Chicago, USA).

### Statistical analysis

Statistical analyses were performed with GraphPad Prism 6 (La Jolla, CA, USA) using two-tailed t-test or one way-Anova with uncorrected Fisher’s LSD. Data was tested for normal distribution with a D’Agostino-Person normality test.

### Ethic statement

*Y. enterocolitica* wild type strain WA-314 was a gift of Jürgen Heesemann (Max von Pettenkofer Institute, Munich, Germany) and described elsewhere [40]. The source of the other strains used here and the generation of *Yersinia* mutants is described in table 1 and in the Methods section.

Approval for the analysis of anonymized blood donations (WF-015/12) was obtained by the Ethical Committee of the Årztekammer Hamburg (Germany).

### Generation of Yersinia mutants

*Y. enterocolitica* mutants WA-314ΔYopB and WA-314ΔYopD were generated as described previously [42], Briefly, mutants were constructed by replacing the coding region of *yopB and yopD* by a kanamycin resistance cassette in the pYV plasmid. Correct replacement of the respective *yop* genes by the resistance cassettes was verified by PCR and SDS-PAGE of secreted Yop proteins and Western blotting. To rule out any unwanted recombination in the chromosome due to the action of Reda and Redp, the mutated plasmids were transferred to the pYV-cured strain WA-C.

### Cell culture and transfection

Hela cells (ACC#57, DSMZ-German Collection of Microorganisms and Cell Cultures) were cultured at 37°C and 5% CO_2_ in DMEM (Invitrogen, GIBCO, Darmstadt, Germany) supplemented with 10% FCS. For infection with bacteria, HeLa cells were seeded in 6 well plates (3×l0^5^ cells per well) or on glass coverslips (12mm, No. 1.5H for high resolution, Marienfeld GmbH, Lauda-Königshafen, Germany) at a density of 5×l0^4^. HeLa cells were transfected with turbofect [Thermo Fisher Scientific, Waltham, Massachusetts, USA) for 8-16h according to the manufacturer’s protocol.

Human peripheral blood monocytes were isolated from heparinized blood as described previously [71]. Monocytes/Macrophages were cultured in RPMU640 [Invitrogen) containing 20% heterologous human serum for 7 days with medium changes every three days. For immunostaining, 1×10^5^ macrophages were seeded on coverslips [12 mm, No. 1.5H, Marienfeld GmbH) one day prior to infection. Macrophages were transfected with the Neon Transfection System [Invitrogen) with 5 μg DNA per 10^6^cells [1000 V, 40 ms, 2 pulses).

### Preparation of bacteria

*Yersinia* were grown in Luria Bertani [LB) broth [supplemented with suitable antibiotics and diaminopimelic acid as stated in Table 1) at 27°C overnight and then diluted 1:20 in fresh broth, followed by cultivation at 37°C for 1.5 h to induce expression of the T3SS. Secretion of Yops was induced by calcium depletion of the *Yersinia* growth medium as described before [72]. Bacteria were centrifuged and resuspended in ice-cold PBS [secretion condition) or in PK solution [500 (μg/ml in PBS, secretion + PK) at RT for 10 min, followed by incubation with 4 mM PMSF in PBS. For immunostaining, bacteria were attached to gelatin [0.2 %) coated coverslips and fixed with 4% para-formaldehyde [PFA; Electron Microscopy Science, Hatfield, USA) for 5 min. Biotinylation of bacteria was performed with EZ-Link™ Sulfo-NHS-SS-Biotin [Thermo Fisher Scientific) as described previously [64]. For cell infection, bacteria were centrifuged, resuspended in ice-cold PBS and added to target cells at a multiplicity of infection [MOI) of 10-100 for 5 to 120 min, as indicated. Target cells in *Yersinia* containing growth medium were centrifuged at 200 x g for 2 min to synchronize the bacterial attachment. Thereafter cells were washed twice with PBS to remove unbound bacteria.

### Immunofluorescence staining

Samples were fixed with 4% PFA in PBS for 5 min and permeabilized with 0.1% Triton X-100 (w/v) in PBS for 10 min. Unspecific binding sites were blocked with 3% bovine serum albumin (BSA, w/v) in PBS for at least 30 min. Bacterial samples were treated with 0.1% Triton X-100 in PBS for non-permeabilizing conditions or with 2% SDS (w/v) in PBS for permeabilizing conditions. Samples were then incubated with a 1:100 dilution of the indicated primary antibody for 1 h, washed three times with PBS and incubated with a 1:200 dilution of the suitable fluorophore-coupled secondary antibody for 45 min. Both, primary and secondary antibodies were applied in PBS supplemented with 3% BSA. Fluorophore-coupled phalloidin (1:200, Invitrogen) and 4’,6-diamidino-2-phenylindole (DAPI; 300 nM, Invitrogen) were added to the secondary antibody staining solution as indicated. Colocalization studies using STED microscopy were performed with Abberior-StarRed and AlexaFluor-594 labelled secondary antibodies. For staining of biotinylated yersiniae Cy5-conjugated streptavidin (strep-Cy5; 1:100, Thermo Fisher Scientific) was added to the primary antibody staining solution. Coverslips were mounted in MOWIOL (Calbiochem, Darmstadt, Germany), ProLond Diamond (Thermo Fisher Scientific) or Abberior mount liquid antifade (Abberior).

### Microscopy and high resolution imaging

Fixed samples were analyzed with confocal laser scanning microscopes (Leica TCS SP5 or SP8) equipped with a 63x, NA1.4 oil immersion objective and Leica LAS AF or LAS X SP8 software (Leica Microsystems, Wetzlar, Germany) were used for acquisition, respectively. STED and corresponding confocal microscopy were carried out in sequential line scanning mode using two Abberior STED setups. The first was based on an Olympus IX microscope body and made use of 100x NA 1.4 oil immersion objective for fluorescence excitation and detection. The second setup, used for colocalization studies, was based on a Nikon Ti-E microscope body with perfect focus system and employed for excitation and detection of the fluorescence signal a 60x (NA 1.4) P-Apo oil immersion objective. Two pulsed lasers were used for excitation at 561 and 640 nm and near-infrared pulsed laser (775 nm) for depletion. The detected fluorescence signal was directed through a variable sized pinhole (set to match 1 Airy at 640 nm) and detected by novel state of the art avalanche photo diodes APDs with appropriate filter settings for Cy3 (605-635 nm) and Cy5 (615-755 nm). Images were recorded with a dwell time of 10 μs and the voxel size was set to be 20×20×150 nm for 2D-STED or 40×40×40 nm for 3D-STED. The acquisitions were carried out in time gating mode i.e. with a time gating width of 8ns and a delay of 781ps (Cy3) and 935ps (Cy5). 3D-STED images were acquired with 80% 3D donut 3D-STED z-stacks were background subtracted and colocalization events were quantified using the ImageJ plugin JACoP. STED spots of 3D-STED images were quantified with Imaris v6.1.1. (Bitplane, Zürich, CH). After baseline subtraction, each channel was analyzed individually with settings adjusted to confocal or STED. Background subtraction was applied in order to detect single spots. An average spot was measured (diameter of largest sphere 0.3-0.45 μm for confocal and 0.11-0.24 μm for STED) and the Gaussian filter was adjusted to the diameter of the largest sphere. Local maxima were filtered by size and quality of spots, which is defined by the intensity at the center of the spot.

### Analysis of clusters formed by YopB/YopD spots visualized with STED

STED spot detection and segmentation was done with Imaris (v6.1.1) on 3D-STED images as described above. Segmentation analysis provided information about volume and position of single STED spots which was then evaluated by a cluster analysis in Matlab (S2 MatLab script, written by AF; Version 9.2, The MathWorks Inc., Natick, USA). In brief, the radius of each spot was calculated through its volume and the resulting sphere was placed at the position of the centre of mass of the segmented volume. The spot size was defined as the full width at half maximum (FWHM) of a fluorescence profile (resolution ~100 nm). Distances between the centers of mass were calculated and categorized into clusters. The threshold for spots included in a cluster was defined by the sum of the radiuses of two spheres. For visualization of STED clusters they were projected in different colors in a superresolution 3D image.

### 3D-SIM (structured illumination microscopy) superresolution microscopy

We used 3D structured illumination microscopy to visualize the GFP-tagged basal body component YscD together with an antibody staining against the pore proteins YopB/D. Cells were imaged with a CFI Apochromat TIRF 100x Oil / NA 1.49 objective (Nikon, Tokio, Japan) on a Nikon N-SIM E equipped with a Ti eclipse inverted microscope (Nikon). Images were acquired using NIS Elements software steering a LU-N3-SIM 488/561/640 laser unit, an Orca flash 4.0 LT (Hamamatsu Photonics, Hamamatsu, Japan) sCMOS or Ultra EM-CCD DU-897 (Andor Technology, Belfast, Northern Ireland) camera, a Piezo z drive (Mad city labs, Wisconsin, USA), a N-SIM motorized quad band filter combined with N-SIM 488 and 561 bandpass emission filters using laser lines 488 and 561 at 100% output power and adjustable exposure times of (300-800 msec). Z stacks were acquired at 200 nm step size, covering about 1.6-2.2 μm. Reconstruction was performed with the stack reconstruction tool (Nikon, NIS-Elements) using default parameters.

### Image analysis of SIM data

Particle detection was done with the ImageJ plugin TrackMate v3.5.1 [73] followed by a nearest neighborhood analysis in Matlab (Version 9.2). In brief, single z-planes of 3D 2-color SIM images were used for analysis. Distinct spots of GFP-YscD and YopB/D were detected with Trackmate using an estimated blob diameter of 0.3 micron for both channels, individually set thresholds and activated median filter. The resulting x/y coordinates were exported into xml files and imported into Matlab by parseXML (github.com/samuellab). The euclidean nearest neighbors of GFP-YscD spots for every YopB/D spot were detected with the function Knnsearch and the resulting distances were plottet Scripts (written by JB) can be found in S2 MatLab script.

### Immunoelectron microscopy

For TEM analysis, Hela cells were transfected with myc-Rac1Q61L for 16 h and infected with E40 GFP-YscD at a ratio of 50:1 for 1 h. Samples were washed with ice-cold PBS and then fixed with a double-strength fixative made of 4% PFA (w/v) with 0.2% glutaraldehyde (GA) in PB. Fixative was replaced after 10 min with fresh single strength fixative (2% PFA with 0.1% GA in PB for 20 min) at RT. Fixation was followed by three wash steps with PBS at RT. Samples were then warmed up at 37°C and coated with 1% gelatin. Sedentary cells were harvested carefully and spinned down in the same fixative. Suspended cells were embedded in 12% (w/v) gelatin in PBS at 37°C for 5 min and the pellet was solidified on ice. Small blocks were cut and infiltrated in 2.3 M sucrose overnight. Thereafter the blocks were mounted on specimen holders and frozen by immersing them in liquid nitrogen. Ultrathin sections (70 nm) were cut and labeled according to Slot and Geuze [58]. Briefly, sections were collected on Carbon-Formvar-coated nickel grids (Science Services GmbH, München, Germany). Anti-GFP-biotin (1:300) was recognized with secondary anti-biotin (1:10 000) and 15 nm large protein A gold; anti-YopD (1:50) with 10 nm large protein A gold in single and double labelling experiments. Sections were examined in an EM902 (Zeiss, Oberkochen, Germany). Pictures were taken with a TRS 2K digital camera (A. Tröndle, Moorenweis, Germany) at 30.000x magnifcation.

Labelling intensities in randomly selected images were quantified as described previously [74, 75]. Briefly, compartments of interest were defined for each antigen (Fig 3E; YscD: bacterial cytoplasm, bacterial double membrane and host cell; YopD: bacterial cytoplasm, injectisome range, host cell). Labelling intensities were then analyzed by counting colloidal gold particles per compartment. Superimposed test-point lattices were used to estimate the compartment area and to count chance encounters with compartments. Labelling densities (gold/μm^2^), expected golds (n_e_) and relative labelling indices (RLI) with partial chi-squared values were then calculated from observed and expected gold distributions to determine the preferentially labelled compartments [75]. For a combined analysis of volume and surface occupying compartments an acceptance zone adjacent to membranes was defined and this zone was treated as a profile area [76].

### Detection of released effector and translocator proteins (Digitonin Lysis Assay)

Released effector (YopH) and translocator proteins (YopB/D) were analyzed as described previously [63]. In brief, HeLa cells were infected with a MOI of 100 for 1 h and subsequently washed with PBS to remove non-adherent bacteria. To remove extracellular effectors, cells were treated with PK (500 mg/ml in PBS) for 20 min at room temperature. Prior to cell lysis, protease activity was blocked by the addition of PMSF (4 mM in PBS). HeLa cells were lysed by the addition of digitonin (0.5% w/v in PBS) at room temperature for 20 min, with repeated vortexing. Cell debris and attached bacteria were separated from the lysate containing the released effectors by centrifugation. The resulting supernatants and pellets were analyzed by SDS-PAGE, transferred to a PVDF membrane (Immobilon-P, Millipore), and analyzed by Western blot using antisera against YopH, YopB and YopD and antibodies against actin, calnexin and myc-tag.

## Acknowledgements

This work is part of the doctoral theses of Franziska Huschka and Theresa Nauth. We thank Frank Bentzien (UKE Transfusion Medicine) for buffy coats, Jürgen Heesemann (formerly Max-von-Pettenkofer Institute, Munich, Germany) for antibodies, bacterial strains and continuous support, Christian Schuberth (Institute of Cell Dynamics and Imaging, University of Münster, Münster, Germany) for excellent support in SIM microscopy, Bernd Zobiak (UKE Microscopy Imaging Facility) for helpful suggestions and support in confocal and STED microscopy and Christian Wurm (Abberior Instruments GmbH, Göttingen, Germany) for providing STED technology and technical support. Further, we are grateful to Nikon Instruments GmbH, Germany, for providing the N-SIM system.

## References

1. Diepold A, Kudryashev M, Delalez NJ, Berry RM, Armitage JP. Composition, formation, and regulation of the cytosolic c-ring, a dynamic component of the type III secretion injectisome. PLoS Biol. 2015;13:e1002039. doi:10.1371/journal.pbio.1002039.

2. Deng W, Marshall NC, Rowland JL, McCoy JM, Worrall LJ, Santos AS, et al. Assembly, structure, function and regulation of type III secretion systems. Nat Rev Microbiol. 2017;15:323–37. doi:10.1038/nrmicro.2017.20.

3. Galán JE, Lara-Tejero M, Marlovits TC, Wagner S. Bacterial type III secretion systems: specialized nanomachines for protein delivery into target cells. Annu Rev Microbiol. 2014;68:415–38. doi:10.1146/annurev-micro-092412-155725.

4. Abby SS, Rocha EPC. The non-flagellar type III secretion system evolved from the bacterial flagellum and diversified into host-cell adapted systems. PLoS Genet. 2012;8:e1002983. doi:10.1371/journal.pgen.1002983.

5. Pinaud L, Sansonetti PJ, Phalipon A. Host Cell Targeting by Enteropathogenic Bacteria T3SS Effectors. Trends Microbiol. 2018;26:266–83. doi:10.1016/j.tim.2018.01.010.

6. Galán JE, Waksman G. Protein-Injection Machines in Bacteria. Cell. 2018;172:1306–18. doi:10.1016/j.cell.2018.01.034.

7. Bai F, Li Z, Umezawa A, Terada N, Jin S. Bacterial type III secretion system as a protein delivery tool for a broad range of biomedical applications. Biotechnol Adv. 2018;36:482–93. doi:10.1016/j.biotechadv.2018.01.016.

8. Anantharajah A, Mingeot-Leclercq M-P, van Bambeke F. Targeting the Type Three Secretion System in Pseudomonas aeruginosa. Trends Pharmacol Sci. 2016;37:734–49. doi:10.1016/j.tips.2016.05.011.

9. Marshall NC, Finlay BB. Targeting the type III secretion system to treat bacterial infections. Expert Opin Ther Targets. 2014;18:137–52. doi:10.1517/14728222.2014.855199.

10. Rüssmann H, Shams H, Poblete F, Fu Y, Galán JE, Donis RO. Delivery of epitopes by the Salmonella type III secretion system for vaccine development. Science. 1998;281:565–8.

11. Dewoody RS, Merritt PM, Marketon MM. Regulation of the Yersinia type III secretion system: traffic control. Front Cell Infect Microbiol. 2013;3:4. doi:10.3389/fcimb.2013.00004.

12. Portaliou AG, Tsolis KC, Loos MS, Zorzini V, Economou A. Type III Secretion: Building and Operating a Remarkable Nanomachine. Trends Biochem Sci. 2016;41:175–89. doi:10.1016/j.tibs.2015.09.005.

13. Kudryashev M, Stenta M, Schmelz S, Amstutz M, Wiesand U, Castaño-Díez D, et al. In situ structural analysis of the Yersinia enterocolitica injectisome. Elife. 2013;2:e00792. doi:10.7554/eLife.00792.

14. Matteï P-J, Faudry E, Job V, Izoré T, Attree I, Dessen A. Membrane targeting and pore formation by the type III secretion system translocon. FEBS J. 2011;278:414–26. doi:10.1111/j.1742-4658.2010.07974.x.

15. Cornelis GR. The type III secretion injectisome. Nat Rev Microbiol. 2006;4:811–25. doi:10.1038/nrmicro1526.

16. Akopyan K, Edgren T, Wang-Edgren H, Rosqvist R, Fahlgren A, Wolf-Watz H, Fallman M. Translocation of surface-localized effectors in type III secretion. Proc Natl Acad Sci U S A. 2011;108:1639–44. doi:10.1073/pnas.1013888108.

17. Mueller CA, Broz P, Cornelis GR. The type III secretion system tip complex and translocon. Mol Microbiol. 2008;68:1085–95. doi:10.1111/j.1365-2958.2008.06237.x.

18. Håkansson S, Schesser K, Persson C, Galyov EE, Rosqvist R, Homblé F, Wolf-Watz H. The YopB protein of Yersinia pseudotuberculosis is essential for the translocation of Yop effector proteins across the target cell plasma membrane and displays a contact-dependent membrane disrupting activity. EMBO J. 1996;15:5812–23.

19. Holmström A, Petterson J, Rosqvist R, Håkansson S, Tafazoli F, Fällman M, et al. YopK of Yersinia pseudotuberculosis controls translocation of Yop effectors across the eukaryotic cell membrane. Mol Microbiol. 1997;24:73–91.

20. Blocker A, Gounon P, Larquet E, Niebuhr K, Cabiaux V, Parsot C, Sansonetti P. The tripartite type III secreton of Shigella flexneri inserts IpaB and IpaC into host membranes. J Cell Biol. 1999;147:683–93.

21. Neyt C, Cornelis GR. Insertion of a Yop translocation pore into the macrophage plasma membrane by Yersinia enterocolitica: requirement for translocators YopB and YopD, but not LcrG. Mol Microbiol. 1999;33:971–81.

22. Dacheux D, Goure J, Chabert J, Usson Y, Attree I. Pore-forming activity of type III system-secreted proteins leads to oncosis of Pseudomonas aeruginosa-infected macrophages. Mol Microbiol. 2001;40:76–85.

23. Miki T, Okada N, Shimada Y, Danbara H. Characterization of Salmonella pathogenicity island 1 type III secretion-dependent hemolytic activity in Salmonella enterica serovar Typhimurium. Microb Pathog. 2004;37:65–72. doi:10.1016/j.micpath.2004.04.006.

24. Ide T, Laarmann S, Greune L, Schillers H, Oberleithner H, Schmidt MA. Characterization of translocation pores inserted into plasma membranes by type III-secreted Esp proteins of enteropathogenic Escherichia coli. Cell Microbiol. 2001;3:669–79.

25. Romano FB, Tang Y, Rossi KC, Monopoli KR, Ross JL, Heuck AP. Type 3 Secretion Translocators Spontaneously Assemble a Hexadecameric Transmembrane Complex. J Biol Chem. 2016;291:6304–15. doi:10.1074/jbc.M115.681031.

26. Montagner C, Arquint C, Cornelis GR. Translocators YopB and YopD from Yersinia enterocolitica form a multimeric integral membrane complex in eukaryotic cell membranes. J Bacteriol. 2011;193:6923–8. doi:10.1128/JB.05555-11.

27. Aepfelbacher M, Roppenser B, Hentschke M, Ruckdeschel K. Activity modulation of the bacterial Rho GAP YopE: an inspiration for the investigation of mammalian Rho GAPs. Eur J Cell Biol. 2011;90:951–4. doi:10.1016/j.ejcb.2010.12.004.

28. Sheahan K-L, Isberg RR. Identification of mammalian proteins that collaborate with type III secretion system function: involvement of a chemokine receptor in supporting translocon activity. MBio. 2015;6:e02023–14. doi:10.1128/mBio.02023-14.

29. Misselwitz B, Dilling S, Vonaesch P, Sacher R, Snijder B, Schlumberger M, et al. RNAi screen of Salmonella invasion shows role of COPI in membrane targeting of cholesterol and Cdc42. Mol Syst Biol. 2011;7:474. doi:10.1038/msb.2011.7.

30. Nichols CD, Casanova JE. Salmonella-directed recruitment of new membrane to invasion foci via the host exocyst complex. Curr Biol. 2010;20:1316–20. doi:10.1016/j.cub.2010.05.065.

31. Viboud GI, Bliska JB. A bacterial type III secretion system inhibits actin polymerization to prevent pore formation in host cell membranes. EMBO J. 2001;20:5373–82. doi:10.1093/emboj/20.19.5373.

32. Mejía E, Bliska JB, Viboud GI. Yersinia controls type III effector delivery into host cells by modulating Rho activity. PLoS Pathog. 2008;4:e3. doi:10.1371/journal.ppat.0040003.

33. Köberle M, Klein-Günther A, Schütz M, Fritz M, Berchtold S, Tolosa E, et al. Yersinia enterocolitica targets cells of the innate and adaptive immune system by injection of Yops in a mouse infection model. PLoS Pathog. 2009;5:e1000551. doi:10.1371/journal.ppat.1000551.

34. Schweer J, Kulkarni D, Kochut A, Pezoldt J, Pisano F, Pils MC, et al. The cytotoxic necrotizing factor of Yersinia pseudotuberculosis (CNFY) enhances inflammation and Yop delivery during infection by activation of Rho GTPases. PLoS Pathog. 2013;9:e1003746. doi:10.1371/journal.ppat.1003746.

35. Wolters M, Boyle EC, Lardong K, Trülzsch K, Steffen A, Rottner K, et al. Cytotoxic necrotizing factor-Y boosts Yersinia effector translocation by activating Rac protein. J Biol Chem. 2013;288:23543–53. doi:10.1074/jbc.M112.448662.

36. Brubaker, R. R., Surgalla, M. J. The effect of Ca++ and Mg++ on lysis, growth, and production of virulence antigens by pasteurella pestis. J Infect Dis. 1964;114:13–25.

37. Michiels T, Wattiau P, Brasseur R, Ruysschaert JM, Cornelis G. Secretion of Yop proteins by Yersiniae. Infect Immun. 1990;58:2840–9.

38. Straley SC, Plano GV, Skrzypek E, Haddix PL, Fields KA. Regulation by Ca2+ in the Yersinia low-Ca2+ response. Mol Microbiol. 1993;8:1005–10.

39. Hell SW, Wichmann J. Breaking the diffraction resolution limit by stimulated emission: stimulated-emission-depletion fluorescence microscopy. Opt Lett. 1994;19:780–2.

40. Heesemann J, Laufs R. Construction of a mobilizable Yersinia enterocolitica virulence plasmid. J Bacteriol. 1983;155:761–7.

41. Oellerich MF, Jacobi CA, Freund S, Niedung K, Bach A, Heesemann J, Trülzsch K. Yersinia enterocolitica infection of mice reveals clonal invasion and abscess formation. Infect Immun. 2007;75:3802–11. doi:10.1128/IAI.00419-07.

42. Trülzsch K, Sporleder T, Igwe EI, Rüssmann H, Heesemann J. Contribution of the major secreted yops of Yersinia enterocolitica O:8 to pathogenicity in the mouse infection model. Infect Immun. 2004;72:5227–34. doi:10.1128/IAI.72.9.5227-5234.2004.

43. Marketon MM, DePaolo RW, DeBord KL, Jabri B, Schneewind O. Plague bacteria target immune cells during infection. Science. 2005;309:1739–41. doi:10.1126/science.1114580.

44. Durand EA, Maldonado-Arocho FJ, Castillo C, Walsh RL, Mecsas J. The presence of professional phagocytes dictates the number of host cells targeted for Yop translocation during infection. Cell Microbiol. 2010;12:1064–82. doi:10.1111/j.1462-5822.2010.01451.x.

45. Pettersson J, Holmström A, Hill J, Leary S, Frithz-Lindsten E, Euler-Matell A von, et al. The V-antigen of Yersinia is surface exposed before target cell contact and involved in virulence protein translocation. Mol Microbiol. 1999;32:961–76.

46. Broz P, Mueller CA, Müller SA, Philippsen A, Sorg I, Engel A, Cornelis GR. Function and molecular architecture of the Yersinia injectisome tip complex. Mol Microbiol. 2007;65:1311–20. doi:10.1111/j.1365-2958.2007.05871.x.

47. Derewenda U, Mateja A, Devedjiev Y, Routzahn KM, Evdokimov AG, Derewenda ZS, Waugh DS. The structure of Yersinia pestis V-antigen, an essential virulence factor and mediator of immunity against plague. Structure. 2004;12:301–6. doi:10.1016/j.str.2004.01.010.

48. Deane JE, Roversi P, Cordes FS, Johnson S, Kenjale R, Daniell S, et al. Molecular model of a type III secretion system needle: Implications for host-cell sensing. Proc Natl Acad Sci U S A. 2006;103:12529–33. doi:10.1073/pnas.0602689103.

49. Johnson S, Roversi P, Espina M, Olive A, Deane JE, Birket S, et al. Self-chaperoning of the type III secretion system needle tip proteins IpaD and BipD. J Biol Chem. 2007;282:4035–44. doi:10.1074/jbc.M607945200.

50. Zhang Y, Lara-Tejero M, Bewersdorf J, Galán JE. Visualization and characterization of individual type III protein secretion machines in live bacteria. Proc Natl Acad Sci U S A. 2017;114:6098–103. doi:10.1073/pnas.1705823114.

51. Hu B, Lara-Tejero M, Kong Q, Galán JE, Liu J. In Situ Molecular Architecture of the Salmonella Type III Secretion Machine. Cell. 2017;168:1065-1074.e10. doi:10.1016/j.cell.2017.02.022.

52. Mueller CA, Broz P, Müller SA, Ringler P, Erne-Brand F, Sorg I, et al. The V-antigen of Yersinia forms a distinct structure at the tip of injectisome needles. Science. 2005;310:674–6. doi:10.1126/science.1118476.

53. Journet L, Agrain C, Broz P, Cornelis GR. The needle length of bacterial injectisomes is determined by a molecular ruler. Science. 2003;302:1757–60. doi:10.1126/science.1091422.

54. Gustafsson MGL, Shao L, Carlton PM, Wang CJR, Golubovskaya IN, Cande WZ, et al. Three-dimensional resolution doubling in wide-field fluorescence microscopy by structured illumination. Biophys J. 2008;94:4957–70. doi:10.1529/biophysj.107.120345.

55. Schermelleh L, Carlton PM, Haase S, Shao L, Winoto L, Kner P, et al. Subdiffraction multicolor imaging of the nuclear periphery with 3D structured illumination microscopy. Science. 2008;320:1332–6. doi:10.1126/science.1156947.

56. Griffiths G. Fine Structure Immunocytochemistry. Berlin, Heidelberg: Springer Berlin Heidelberg; 1993.

57. Koster AJ, Klumperman J. Electron microscopy in cell biology: integrating structure and function. Nat Rev Mol Cell Biol. 2003;Suppl:SS6–10.

58. Slot JW, Geuze HJ. Cryosectioning and immunolabeling. Nat Protoc. 2007;2:2480–91. doi:10.1038/nprot.2007.365.

59. Aili M, Isaksson EL, Carlsson SE, Wolf-Watz H, Rosqvist R, Francis MS. Regulation of Yersinia Yop-effector delivery by translocated YopE. Int J Med Microbiol. 2008;298:183–92. doi:10.1016/j.ijmm.2007.04.007.

60. Aepfelbacher M, Wolters M. Acting on Actin: Rac and Rho Played by Yersinia. Curr Top Microbiol Immunol. 2017;399:201–20. doi:10.1007/82_2016_33.

61. Black DS, Bliska JB. The RhoGAP activity of the Yersinia pseudotuberculosis cytotoxin YopE is required for antiphagocytic function and virulence. Mol Microbiol. 2000;37:515–27.

62. Pawel-Rammingen U von, Telepnev MV, Schmidt G, Aktories K, Wolf-Watz H, Rosqvist R. GAP activity of the Yersinia YopE cytotoxin specifically targets the Rho pathway: a mechanism for disruption of actin microfilament structure. Mol Microbiol. 2000;36:737–48.

63. Nordfelth R, Wolf-Watz H. YopB of Yersinia enterocolitica is essential for YopE translocation. Infect Immun. 2001;69:3516–8. doi:10.1128/IAI.69.5.3516-3518.2001.

64. Sarantis H, Balkin DM, Camilli P de, Isberg RR, Brumell JH, Grinstein S. Yersinia entry into host cells requires Rab5-dependent dephosphorylation of PI(4,5)P_2_ and membrane scission. Cell Host Microbe. 2012;11:117–28. doi:10.1016/j.chom.2012.01.010.

65. Bahnan W, Boettner DR, Westermark L, Fällman M, Schesser K. Pathogenic Yersinia Promotes Its Survival by Creating an Acidic Fluid-Accessible Compartment on the Macrophage Surface. PLoS ONE. 2015;10:e0133298. doi:10.1371/journal.pone.0133298.

66. Wong K-W, Mohammadi S, Isberg RR. The polybasic region of Rac1 modulates bacterial uptake independently of self-association and membrane targeting. J Biol Chem. 2008;283:35954–65. doi:10.1074/jbc.M804717200.

67. Alrutz MA, Srivastava A, Wong KW, D’Souza-Schorey C, Tang M, Ch’Ng LE, et al. Efficient uptake of Yersinia pseudotuberculosis via integrin receptors involves a Rac1-Arp 2/3 pathway that bypasses N-WASP function. Mol Microbiol. 2001;42:689–703.

68. Wong K-W, Isberg RR. Arf6 and phosphoinositol-4-phosphate-5-kinase activities permit bypass of the Rac1 requirement for beta1 integrin-mediated bacterial uptake. J Exp Med. 2003;198:603–14. doi:10.1084/jem.20021363.

69. Liu J, Park D, Lara-Tejero M, Galan JE, Li W, Waxham MN, Hu B. Visualization of the type III secretion mediated Salmonella-host cell interface using cryo-electron tomography; 2018.

70. Schmid A, Neumayer W, Trülzsch K, Israel L, Imhof A, Roessle M, et al. Cross-talk between type three secretion system and metabolism in Yersinia. J Biol Chem. 2009;284:12165–77. doi:10.1074/jbc.M900773200.

71. Linder S, Nelson D, Weiss M, Aepfelbacher M. Wiskott-Aldrich syndrome protein regulates podosomes in primary human macrophages. Proc Natl Acad Sci U S A. 1999;96:9648–53.

72. Heesemann J, Gross U, Schmidt N, Laufs R. Immunochemical analysis of plasmid-encoded proteins released by enteropathogenic Yersinia sp. grown in calcium-deficient media. Infect Immun. 1986;54:561–7.

73. Tinevez J-Y, Perry N, Schindelin J, Hoopes GM, Reynolds GD, Laplantine E, et al. TrackMate: An open and extensible platform for single-particle tracking. Methods. 2017;115:80–90. doi:10.1016/j.ymeth.2016.09.016.

74. Griffiths G, Hoppeler H. Quantitation in immunocytochemistry: correlation of immunogold labeling to absolute number of membrane antigens. J Histochem Cytochem. 1986;34:1389–98. doi:10.1177/34.11.3534077.

75. Mayhew TM, Lucocq JM, Griffiths G. Relative labelling index: a novel stereological approach to test for non-random immunogold labelling of organelles and membranes on transmission electron microscopy thin sections. J Microsc. 2002;205:153–64.

76. Mayhew TM, Lucocq JM. Quantifying immunogold labelling patterns of cellular compartments when they comprise mixtures of membranes (surface-occupying) and organelles (volume-occupying). Histochem Cell Biol. 2008;129:367–78. doi:10.1007/s00418-007-0375-6.

